# Not Just Noise: Impaired Oscillatory Entrainment Reflects Reduced Temporal Flexibility in Autism

**DOI:** 10.1101/2025.07.02.662876

**Authors:** Shlomit Beker, Theo Vanneau, Elizabeth Akinyemi, John J. Foxe, Sophie Molholm

## Abstract

Rhythmic patterns in the environment enhance neural activity, perception, and action. However, natural rhythms are often imprecise, requiring flexible adaptation. In autism Spectrum Disorder (ASD), characterized by cognitive rigidity and atypical use of prior information - favoring immediate sensory input over predictive cues - entrainment to temporally variable input may be reduced at both neural and behavioral levels, though the neural mechanisms remain unclear. Here, we recorded high-density EEG and behavior in adults with ASD (n=20) and neurotypical (NT) controls (n=21) during a visual detection task with four rhythmic structures, parametrically varied from an isochronous fully regular rhythm, to a highly irregular one. Spectral analysis and temporal response function (TRF) models revealed significantly reduced modulation by temporal regularity in ASD, particularly in mildly jittered stimulation streams. Additionally, the coupling between phases of neural oscillations and behavior was diminished in ASD under the jittered conditions, suggesting reduced functional relevance of neural synchronization. Residual spectral power post-stimulation showed lower oscillatory entrainment in ASD, ruling out simple evoked-response explanations. Notably, the degree of neural modulation by temporal regularity was correlated with IQ within the ASD group, suggesting a link between temporal flexibility and individual cognitive profiles. These findings highlight impaired neural entrainment and reduced behavioral modulation by temporal structure in ASD, offering insight into inflexible responses to uncertain, volatile sensory environments.

**Innovation:** Entrainment to rhythmic events is reduced in autism, but it remains unclear whether this reflects a general, non-selective deficit in neuro-oscillatory alignment or a selective vulnerability to volatile temporal structures, such as those with embedded jitter. To address this, we recorded cortical activity and behavioral performance as participants with ASD engaged with visual sequences of varying rhythmic regularity, and examined how temporal predictability modulated oscillatory entrainment. By correlating neural entrainment with target detection and clinical profiles, we sought to uncover a key feature of the autistic phenotype: reduced temporal flexibility in adapting to unpredictable sensory environments.

## Introduction

Events in natural environments tend to have temporal structure. The nervous system is highly sensitive to these structures (Lakatos *et al*., 2008; Schroeder & Lakatos, 2009; Gomez-Ramirez *et al*., 2011; Wilson & Foxe, 2020; Charalambous & Djebbara, 2023), and leverages the temporal information to optimize perception and action, which in turn facilitates interactions with the surroundings (Rohenkohl *et al*., 2012; Fitch, 2013; Large & Gray, 2015). Stimuli such as natural sounds, body movement and speech, all have rhythmic structures where the occurrence of one event provides a strong predictor of subsequent events (Hutchinson & Barrett, 2019).

This leads to a preference for stimuli that align with expected temporal windows and results in enhanced perceptual performance for those stimuli (Cui *et al*., 2009; Kelly *et al*., 2009; Rohenkohl *et al*., 2012; Cravo *et al*., 2013; Gray *et al*., 2015; Mercier *et al*., 2015). Processing of such events is enhanced through the alignment of ongoing neuro-oscillatory activity to their temporal structure, also referred to as neural entrainment (Lakatos *et al*., 2008; Schroeder *et al*., 2010; Calderone *et al*., 2014; Kosem *et al*., 2014; Arnal & Kleinschmidt, 2017). Notably, neural entrainment occurs even when rhythmic events contain inherent temporal variability (e.g., speech (Ding *et al*., 2016; Luo & Ding, 2024) or are presented in noisy or otherwise suboptimal conditions (Tal *et al*., 2017; Ten Oever *et al*., 2017). This indicates a remarkable tolerance of the nervous system to tune in to imperfect or “noisy” temporal regularities and suggests that neural entrainment represents a significant mechanism by which the nervous system optimizes processing of regularities in the natural, inherently noisy environment. Emerging evidence shows that such entrainment mechanisms might be impaired in ASD which could contribute to the autism phenotype.

Individuals with autism spectrum disorder (ASD) have impaired social communication skills and exhibit rigidity of routines and restricted and highly focused interests (Happe & Frith, 2006). Studies from recent years show reduced adaptation to changes in the environment, and more specifically, compromised tempo-keeping and reduced synchronization to rhythmic stimuli (Lieder *et al*., 2019; Cannon *et al*., 2021; Vishne *et al*., 2021; Kasten *et al*., 2023) (but see (Cannon *et al*., 2023)). However, a characterization of rhythmic event processing, and the conditions under which these impairments emerge in ASD, are limited. Reduced entrainment to rhythmic events may be attributed to different explanations. One possibility is that it results from an overall elevated level of neural noise inherent to the disorder (Rubenstein & Merzenich, 2003; Magnuson *et al*., 2020). Some studies suggest that individuals with ASD exhibit higher variability in both brain activity and behavioral performance compared to controls (Dinstein *et al*., 2012; Beker *et al*., 2021b), although others suggest otherwise (Butler *et al*., 2017; Hecker *et al*., 2022). In this case, entrainment to both isochronous rhythms and rhythms with jittered regularities should be equally impaired.

Otherwise, altered entrainment to rhythmic events could stem from a reduced tolerance of the nervous system to imperfections within a temporal structure—potentially due to hyper-accurate neural representations in ASD (O’Riordan *et al*., 2001; Bonnel *et al*., 2003b; Happe & Frith, 2006; Robertson *et al*., 2013). Under this account, neural representations of temporal events in noisy conditions would be exceptionally impaired, leading to a greater deficit in processing rhythms with jitter compared to fully regular, isochronous rhythms.

We assessed these competing accounts here by systematically manipulating the degree of temporal regularity in sequences of simple visual stimuli. Adult participants diagnosed with autism and neurologically typical (NT) controls viewed sequences of visual stimuli and performed a target detection task. Regularity of the stimuli was manipulated by adding jitter to a fixed stimulus-onset-asynchrony (SOA) in four levels, as follows: *Isochronous* (fixed SOA), and three jitter conditions: *Small Jitter*, *Moderate Jitter* and *Large Jitter*, where SOAs were taken from windows with increasing sizes of jitter. Given that rhythmic events in natural environments are usually not perfectly periodic, a certain level of tolerance for temporal jitter must be maintained for entrainment to be flexible and efficient. A system lacking such tolerance would be expected to not adapt to stimuli flexibly. On the other hand, if the system is too tolerant to jitter, it would not distinguish between useful temporal information and random, unstable, variance. Our analysis was designed to investigate whether neural entrainment and target detection are uniformly reduced in the autism group for all conditions or rather are selectively impaired when jitter is introduced to the regular rhythm, i.e., where regularity exists, but is masked by a jitter.

Coupling of neural oscillations to sensory stimuli has been shown to strongly influence task performance (Fiebelkorn *et al*., 2011; Fiebelkorn *et al*., 2013; Aussel *et al*., 2023). Such oscillations can be stimulus-driven and measured at the frequency of stimulation, or can arise endogenously. Whether and under which conditions oscillatory activity in autism affects performance has not been systematically tested. Here we test the coupling between phase and performance, with the hypothesis that reduced attunement to rhythmic stimuli with jitter results in reduced coupling between phases and reaction times, i.e, performance will show a weaker “preference” for certain phases. We also test if this coupling appears exclusively for the stimulus-driven slow oscillations, or exists in other endogenous oscillations in theta, associated with memory and cognitive control, and alpha, associated with regulation of neural excitability.

Next, we trained a Temporal Response Function (TRF) model to predict EEG activity based on stimulus input. With this we aimed to examine the extent to which neural responses track rhythmic structures with different levels of jitter, on the single-epoch level. TRF models estimate the mapping function between stimulus attributes and resulting brain activity (Lalor *et al*., 2006; Lalor & Foxe, 2010; Crosse *et al*., 2021). By providing a linear approximation of how sensory input is reflected in EEG signals, this approach allows us to quantify neural entrainment by assessing the model’s predictive accuracy: the stronger the alignment between slow EEG fluctuations (0.5–3 Hz) and the presented stimuli, the better the model’s performance. A key advantage of TRF modeling, particularly in clinical populations, is its ability to generate participant-specific models, thereby accounting for the high variability in sensory processing that often characterizes ASD (Schauder & Bennetto, 2016; Delgado-Lobete *et al*., 2020). By training subject-dependent models on EEG data recorded during each condition, we aimed to determine whether neural entrainment in ASD is generally reduced across all levels of temporal regularity or selectively impaired in the presence of jitter. If ASD is associated with a global reduction in neural entrainment, the model’s prediction accuracy should be lower across all conditions. Alternatively, if individuals with ASD exhibit specific difficulty processing irregular rhythms, model performance should be most impaired in the jittered conditions.

Finally, it has been argued that rhythmic entrainment could reflect a sequence of evoked responses (Zoefel *et al*., 2017; van Bree *et al*., 2021; Menétrey & Pascucci, 2024) rather than endogenous entrainment of neural oscillations to patterns of stimulation. To address this issue, we took a two-pronged approach. First, we measured the presence of low-frequency oscillations following the offset of each of the stimulus sequences, when stimulation was no longer present. Resonant low frequency oscillations at the stimulation frequency, in the absence of stimulation, would lend support to an entrainment explanation. An additional approach was to test whether group differences were present in the sensory evoked response. Finding that the VEP does not differ in ASD whereas oscillatory entrainment does (as in (Beker *et al*., 2021b)) would lend support to the idea that the entrainment effect is unrelated to immediate evoked responses.

## Material and Methods

### Participants

(see Table S1 for demographics). Forty-seven individuals participated in the experiment: 23 diagnosed with ASD and 24 age- and IQ-matched neurotypical (NT) controls. Data from three ASD and three NT participants were excluded for poor behavioral performance or insufficient EEG data (see below), retaining 21 NT and 20 ASD participants aged 18–37 (Mean±SD: NT = 21.17 ± 5.8; ASD = 22.5 ± 3.4). IQ scores (PIQ, VIQ, FSIQ) showed no group differences (e.g., FSIQ: NT = 108 ± 13; ASD = 100 ± 16; p = 0.11). ASD diagnosis required: (1) ADOS-2; (2) DSM-5 criteria; and (3) clinical evaluation by an experienced clinician. NT participants had no history of neurological, developmental, or psychiatric disorders, nor first-degree relatives with ASD. All participants (or guardians) gave informed consent under protocols approved by the Albert Einstein College of Medicine institutional review board (IRB). Participants were compensated $15/hour for their participation.

### Stimuli and Task

(See Figure 1 for illustration): The experiment was programmed in Presentation® (Neurobehavioral Systems). Stimuli consisted of grayscale checkerboards (5.9° × 5.9°), composed of 10×10 diagonally divided squares (light gray: 43.3 cd/m^2^; dark gray: 8.3 cd/m^2^; background: 25.2 cd/m^2^). During each condition block, the checkerboard rotated 90° left or right with varying stimulus-onset asynchronies (SOAs). Participants monitored the checkerboard for a target stimulus - a transient, suprathreshold contrast increase - and responded via mouse click. Targets occurred randomly every 8-12 stimuli (6 per block, on average). Each participant’s target contrast was calibrated using an up-down transformed response (UDTR) procedure (Zwislocki & Relkin, 2001) to yield ∼75% detection threshold. The experiment comprised seven SOA conditions (average rate: 1.5 Hz; mean SOA: ∼666 ms); four are reported here. Temporal jitter was introduced to manipulate stimulus regularity by randomly selecting SOAs from uniform distributions (generated via MATLAB function: `unifrnd`) to alter the 666 ms base interval), with increasingly wider bounds: Isochronous (no jitter), Small Jitter (±111 ms), Moderate Jitter (±333 ms), and Large Jitter (±500 ms). Three additional conditions (not reported here) included linearly increasing SOAs, alternating SOAs, and a randomized jitter sequence. Participants completed up to 8 sessions (21 blocks each, 3 blocks per condition), to a total of ∼24 blocks per condition (exceptions: one NT and two ASD completed 9 sessions; one NT completed 7; and one NT completed 5). Short blocks and frequent breaks were used to maintain attention. The full session lasted ∼3 hours, including consent, preparation, and EEG recording.

**Figure 1:**
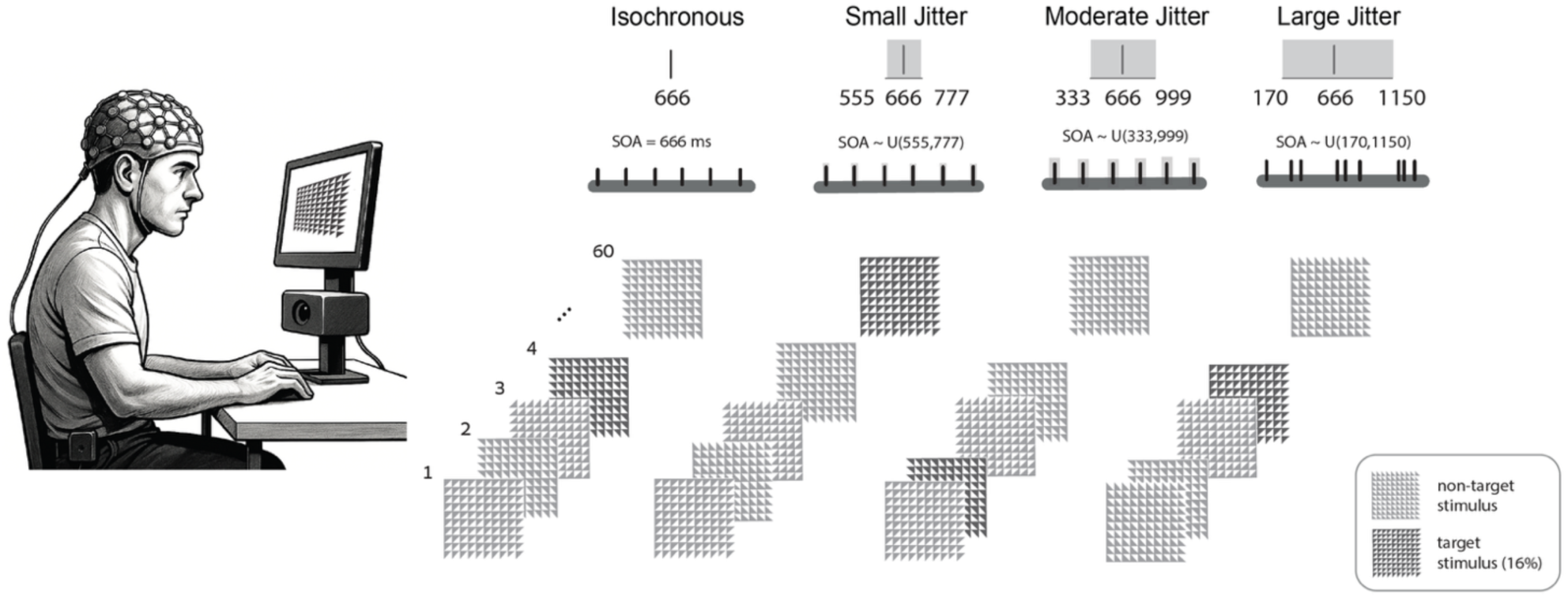
Illustration of experimental design. A visual task is presented to participants while they are being recorded with 64-channel EEG. Stimuli are sequences of 60 checkerboards rotating at different rhythms. From left to right: *Isochronous*, *Small Jitter*, *Moderate Jitter*, and *Large Jitter* conditions, including SOA sizes. Participants are instructed to click the mouse’s button as soon as they detect the occasional higher-contrast targets (in a dark shades).

### Data acquisition

EEG was recorded using BioSemi ActiveTwo with an anti-aliasing filter (-3dB at 3.6 kHz; 70 electrodes: 64 scalp, 6 external) at 512 Hz. Electrodes positions followed the 10-20 system; external electrodes included bilateral mastoids and vertical EOG. Analog triggers for stimulus onsets and button presses were sent from Presentation® and digitized with the EEG stream. Reaction times were recorded via Presentation® software.

### Data and Statistical Analysis

Unless stated otherwise, repeated-measures ANOVAs were used to test Group, Condition, and interaction effects.

### Performance

*Reaction times*: Participants responded to high-contrast checkerboard targets. Valid reaction times (*‘Hits’*) were defined as responses within 150–650 ms post-target. This range excluded anticipatory and late responses. Responses outside this window were labeled false alarms.

#### EEG

Data were processed using MATLAB (R2023a) and FieldTrip (Oostenveld *et al*., 2011). EEG was downsampled to 256 Hz and bandpass-filtered between 0.1–55 Hz using a 5th-order Butterworth IIR filter. Epoching was performed on filtered data. A two-stage artifact rejection procedure was applied at the single-trial level. First, bad channels were identified as those deviating by >1 SD from the mean voltage or auto-covariance across channels; trials with more than six such channels were excluded. Interpolation for acceptable trials used nearest-neighbor spline (Perrin *et al*., 1987). Second, a ±80 μV threshold was used to exclude remaining artifacts. All epochs were demeaned and baseline corrected. Unless mentioned otherwise, measures from occipital channels (O1, Oz, O2, Iz) are reported. Sub-pipelines addressed specific analyses:

### Visual Evoked Response (VEP)

Data were epoched from −200 to +400 ms around each stimulus and baseline-corrected using the −100 to 0 ms window. Signals were referenced to AFz to optimize occipital VEP visualization. Group differences were evaluated by comparing maximal values of P1 peak (140 ± 20 ms) across these channels, with running t-test. Cluster-based permutation tests (2000 permutations) were used to identify significant spatiotemporal differences.

<u>Spectral analysis</u>: was performed using two pipelines: *<u>(1) Power S</u>*<u>pectrum Analysis</u>: Power at 1.5 Hz was computed using Welch’s method (pwelch) with an overlap of 90% to increase spectral resolution, on 5.5-second EEG segments starting 0.5 sec before the first stimulus in each block (covering ∼8 stimuli). A two-way ANOVA tested Group and Condition effects on 1.5 Hz power. *<u>(2) Phase Coherence (ITPC):</u>* ITPC was computed using the formula from (Luo & Poeppel, 2007): 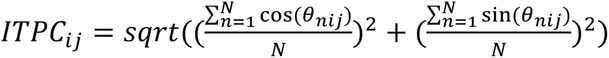, where *θ_nij_* was extracted per trial, frequency, and time point. Values ranged from 0 (no phase alignment) to 1 (perfect alignment). Epochs spanned −1 to +1.4 sec relative to each stimulus. While epochs partially overlapped, the post-stimulus window of interest (∼300 ms) remained uncontaminated. To examine entrainment strength, we analyzed ITPC across the full frequency range (0.6–5 Hz), averaging across time. For group comparisons, we focused on peak ITPC within 1.4–1.8 Hz — the range where most participants showed maximum phase coherence. A two-way repeated measure ANOVA (Group × Condition) tested differences in ITPC amplitude. This was used for the following: <u>ITPC Modulation by Rhythmic Regularity:</u> We quantified ITPC sensitivity to condition by regressing each participant’s ITPC values (1.4–1.6 Hz) against regularity level (4 conditions). Slopes were estimated using the Theil-Sen Estimator (Sen, 1968), a robust non-parametric method suited for four monotonic points. Channel-level slopes were averaged across five predefined clusters: occipital, parieto-occipital, parietal, central, and frontal. Between-group differences were tested using Wilcoxon rank-sum tests. For visualization, Spearman correlations between condition and ITPC were computed at the group level for each region. <u>Time–Frequency Decomposition</u> (Figure S5): Morlet wavelets ((1–20 Hz in 30 steps) with width scaled from 3 cycles at 1 Hz to 5 cycles at 20 Hz were used. Decomposition was performed from −1 to +1.4 sec relative to each stimulus.

### Phase-RT coupling

Participants were included in this analysis if they had at least 60 correct target responses (Hits) in each condition, resulting in exclusion of 2 NT and 1 ASD participants, which resulted in N=19 per group. EEG data were low-pass filtered at 1.5 Hz, and for each target trial, the phase of delta oscillations (1.4–1.6 Hz) at target onset and the corresponding reaction time (RT) were extracted. RTs were binned into eight phase bins (centered: −*π*, −3/4*π*, −*π*/2, −*π*/4, 0, *π*/4, *π*/2, 3/4*π*), and average RTs were computed per bin, per participant. A one-cycle sinusoid was then fit to each participant’s 8-point RT-by-phase curve using a 3-parameter model (amplitude, phase shift, and offset), optimized with MATLAB’s `fminsearch` function. The outcome measure was the peak-to-trough amplitude of this fitted curve. To generate a control distribution, RT-phase assignments were shuffled 1000 times per participant, preserving the RT values but randomly reassigning their bin positions. Sinusoidal fits and amplitude extraction were repeated for each permutation, and average amplitude across iterations was used for comparison. A permutation test was then used to compare real vs. shuffled data across groups and conditions. To assess frequency specificity, the same procedure was repeated at two control frequency bands: theta (5–7 Hz) and alpha (9–11 Hz).

### Resting State Entrainment

The analysis approach was similar to the ITPC elaborated above: We applied ITPC on 10-second EEG epochs that immediately followed each condition block, starting at 2 second following the last stimuli offset, to avoid the VEP effects

### TRF model

We implemented forward encoding models using the `mTRFpy` toolbox (Bialas *et al*., 2023) to assess how well the timing of stimuli could predict neural responses across conditions and groups. EEG data were filtered between 0.1–30 Hz and epoched from −5 to +40 seconds relative to block onset. Artifact rejection followed two stages: first, bad channels were identified and rejected using the `autoreject` toolbox (Jas *et al*., 2017), followed by ICA to remove ocular artifacts (e.g., blinks, saccades). Trials with persistent artifacts were removed, averaging 2.4 ± 0.6 trials for NT and 3.1 ± 0.8 for ASD. Each stimulus stream was encoded as a binary time series reflecting onset timing. EEG was further filtered (0.5–3 Hz) to isolate low-frequency entrained activity. Per subject and condition, 75% of the epochs were randomly assigned to training and 25% to testing (each subject had 24 epochs per condition on average). We refined the model through ridge regression, employing leave-one-out cross-validation, and selecting from the following set of values:

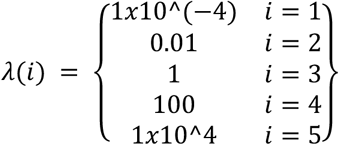

Model performance was quantified by correlating predicted and actual EEG (Pearson’s r) within a 500–2500 ms post-stimulus window to avoid contamination from initial evoked responses. Analysis focused on occipital electrodes (PO3, O1, POz, Oz, Iz, PO4, O2), where visual entrainment was strongest.

### Clinical – Physiological Relationship

Within-group Pearson correlations (FDR-corrected (Benjamini & Hochberg, 1995) were computed between clinical and physiological measures. Clinical scores included in the ASD group: SRS, AQ, FSIQ, ADOS and RBS-R. In NT: SRS, AQ, and FSIQ. Physiological measures included: FFT peak power at 1.5Hz, VEP amplitude, TRF prediction accuracy, target detection accuracy rate and ITPC modulation slope.

## Results

### Behavioral Results

Hit rates: Hit rates (*hit/(hit+misses)*) were higher for the NT group (*mean ± sem, NT: 0.74 ± 0.04; ASD: 0.68 ± 0.04; ranksum = 57519; p = 0.0002*).

*Reaction times*: across all conditions, ASD participants responded slower than controls (F = 499; df = 1; p < 0.001), (RT per condition: NT: 297 *± 10* ms; 276 *± 7* ms; 283 *± 7* ms; 279 *±* 7 ms; ASD: 402+5 ms; 395+5 ms; 401+5 ms; 400+5 ms, for conditions *Isochronous, Small Jitter, Moderate Jitter, Large Jitter*, respectively). No significant effect for Condition (F = 1.11; df = 3; p = 0.34) or Group x Condition interaction (F = 0.31; df = 3; p = 0.8). See Figure 4A.

### Visual Evoked Potentials

The checkerboard stimuli elicited clear visual evoked potentials (VEP) (See Figure 2B) that were highly similar across conditions and between groups, and resembled the VEP elicited by this type of stimulus in prior work (Gray *et al*., 2015; Wilson & Foxe, 2020). A small negative-going response peaked at about 100 ms was followed by a large positive-going response that peaked at about 140 ms. The amplitude of the P1 (peak positive response between 120-160 ms) did not differ as a function of Group or Condition (*Mean ± SEM across conditions, NT: 2.18 ± 0.43 µV; ASD: 1.99 ± 0.34 µV;* Group: *F=0.45; d=1; p=0.5*; Condition: *F=0.69; d=3; p=0.6*; Group × Condition: *F=0.19; d=3; p=0.9)*. The lack of group differences in the VEP is corroborated by the SCP analyses in which all timepoints and channels are considered (Figure 2B). Similarly, inter-trial variability (ITV) of VEP amplitude was not significantly different for any of these factors (Group: *F=3.21; d=1; p=0.07*; Condition: *F=0.15; d=3; p=0.93*; Group × Condition: *F=0.28; d=3; p=0.83)*.

**Figure 2:**
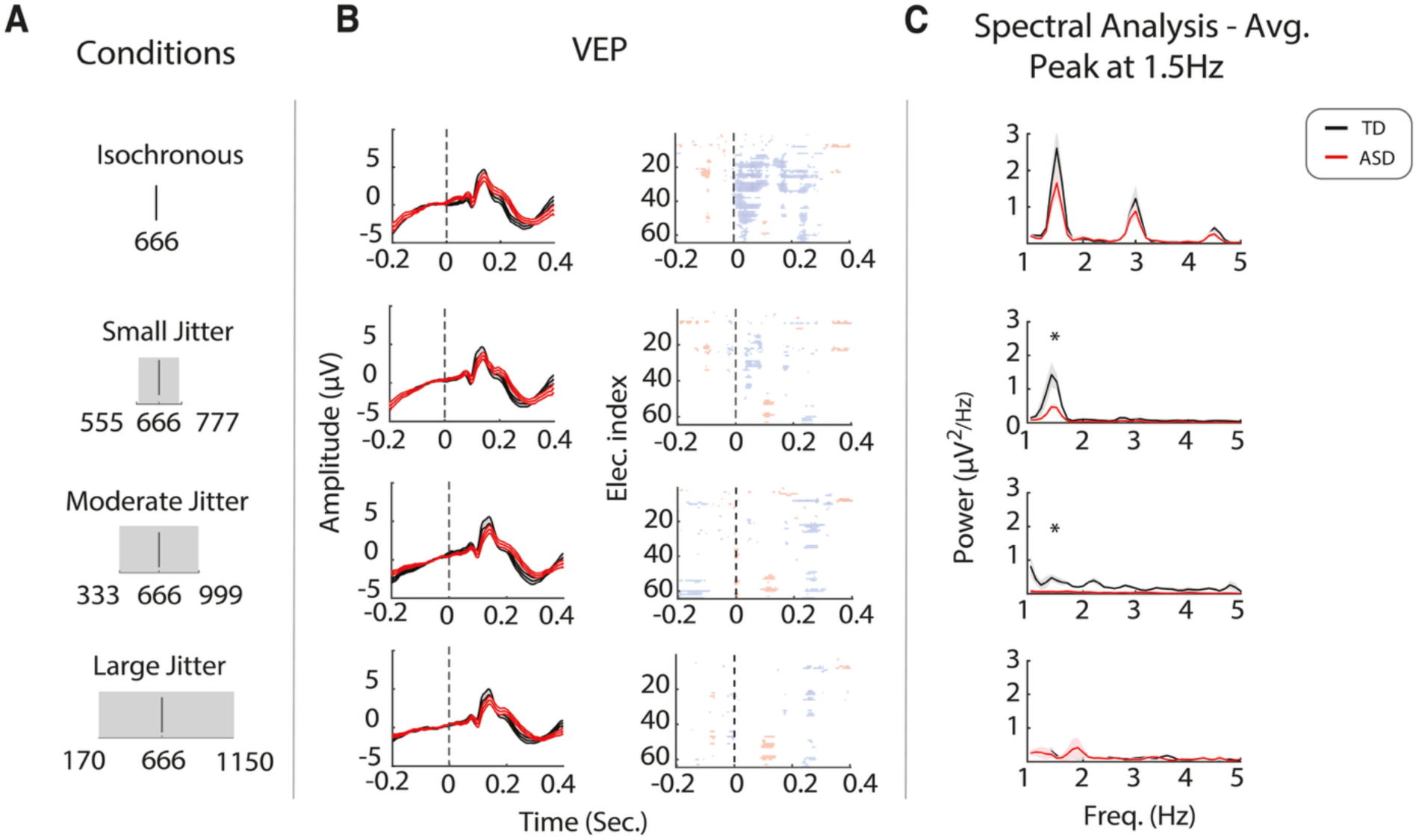
Time- and frequency spectra analysis results show selective group effect in frequency power, but not amplitude, by the conditions. **A.** Condition schematic. **B.** Visual evoked potential (VEP) in each condition, for NT and ASD (left column) and statistical cluster plots (right column) comparing the amplitude of the VEP between groups across all channels and timepoints. No significant difference survived false discovery rate permutation test. Waveforms show average voltage recorded from occipital and parietal channels, where the VEP was strongest. Statistical plots show all channels. **C.** Spectral analysis for group differences in each condition. Anova shows no Group*Condition interaction, however individual tests in each condition show significant group differences in Small and Moderate Jitter conditions (2^nd^ and 3^rd^ rows) in power at 1.5 Hz.

### Spectral analysis

As expected, both groups show a gradual reduction of maximal power at the frequency of stimulation (1.5Hz) with each reduction of periodicity (*NT: mean±SEM: 1.11±0.3 µV^2^; 0.68±0.2 µV^2^; 0.29±0.1 µV^2^; 0.16±0.07 µV^2^. ASD: 0.66±0.19 µV^2^; 0.23±0.06 µV^2^; 0.04±0.01 µV^2^; 0.13±0.09 µV^2^*, for *Isochronous*, *Small*, *Moderate and large Jitter*, respectively). Main effects were found for Group (F = 6.14; df=1; p = 0.01) and Condition (F = 8.8; df = 3; p<0.001), whereas the Group x Condition interaction was not significant (F = 0.75; df = 3; p=0.52). Our initial hypothesis, that a lower modulation of 1.5Hz stimuli will be worse in ASD when presented with jittered regularities, was not supported by the omnibus ANOVA. However, since traditional ANOVA lacks sensitivity to modest effects that are localized to specific conditions, we conducted one-sided permutation tests in each of the four conditions, as a complementary approach. Following Bonferroni correction for multiple comparison, differences were present in the *Small Jitter* (Δ = 0.59 *µV^2^*; Cohen’s d = 0.81; p = 0.028); and *Moderate Jitter* (Δ = 0.29 *µV^2^*; Cohen’s d = 1.15; p = 0.004) conditions, but not for *Isochronous* (Δ = 0.46 *µV^2^*; d = 0.36; p = 0.16) or *Large Jitter* (Δ = 0.03 *µV^2^*; d = 0.09; p = 0.41) conditions. This supports the hypothesis of condition-dependent effects.

Inter-trial phase coherence (ITPC) at the stimulation frequency (1.5 Hz), shown as topographical plots in Figure 3A, gradually decreased as stimulus periodicity decreased—that is, less periodic stimuli led to reduced ITPC. Furthermore, the ASD group showed reduced ITPC relative to controls, for every cortical area of interest. We calculated the Pearson correlation coefficient across individuals in each group (Figure 3B,3C), for visualization purposes. A linear regression fit with Theil-Sen Estimator was calculated for each participant between the average ITPC and the 4 conditions. We found significantly different slopes for ASD than NT for all scalp regions of interest except the frontal region (*NT: mean±SEM: -0.08±0.03, -0.07±0.03, -0.07±0.03, - 0.05±0.02, -0.02±0.01, -0.05±0.02*, for regions: Occipital, Parieto-occipital, Parietal, Central, and Frontal, respectively. *ASD: mean±SD: -0.05±0.03, -0.04±0.03, -0.04±0.03, -0.03±0.02, -0.02±0.01, -0.03±0.02*, for the same scalp regions; Figure 3D), indicating a lower gradation of change in oscillatory synchrony as regularity decreases, at most cortical areas.

**Figure 3:**
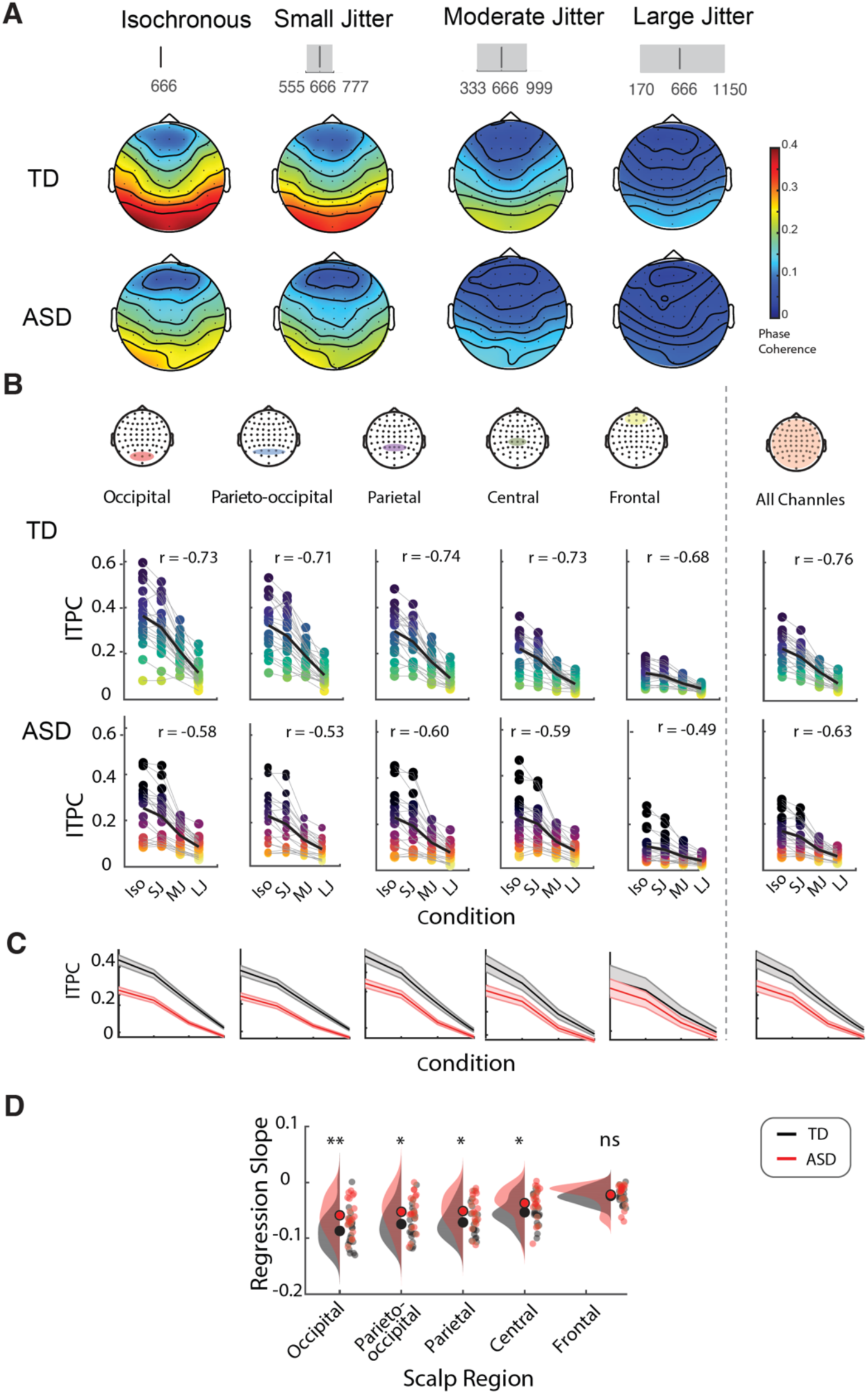
Reduced modulation of inter-trial phase coherence (ITPC) by periodicity is found for ASD group. **A.** ITPC topography plots show the gradated pattern across cortical channels for NT and ASD. **B.** ITPC values in each condition presented for different cortical regions. Dots represent the individuals **C.** Group average slopes. **D.** Rain-cloud plots for individual slopes in each cortical region. All cortical areas but frontal show significantly reduced (shallower) slopes for ASD than NT. (**p<0.01; *p<0.05).

### RT – phase coupling

To test for a relationship between phase of oscillation with RT, we calculated the phase of oscillations at the mean stimulation frequency of 1.5 Hz (see methods) at the time of target onset and measured it as the peak-to-trough amplitude of a one-cycle sinusoid fit.

As mentioned before (*Behavioral Results*), the number of *Hit* trials per participant did not differ between the groups. The distribution of Hit proportion across phases was similar overall in the two groups, with a higher Hit response rate at optimal phase (0) and gradual decrease for phases further away from it for all conditions (Figure 4B). When comparing effects of Group and Condition on the two extreme conditions (*Isochronous* and *Large Jitter*), we found a main effect for Group (p = 0.007), and marginal effect of Condition (p = 0.05), supporting the higher modulation of reaction time by phase in the NT group, with a small advantage for regular vs. irregular rhythm (Figure 4C). We analyzed RT-Phase coupling as peak-to-trough amplitude of a 1-cycle sine fit (Figure 4D-F). Permutation tests, with shuffled data, revealed significant coupling for all conditions in the NT group. In the ASD group, significant coupling was found only in the *Isochronous* condition (Figure 4E). A direct comparison of groups and conditions found main effects for Group (F=15.9; df = 1; p = 0.0001) but not for Condition (Condition: F = 0.32; df = 3; p = 0.81;) or Group × Condition interaction (F = 0.05; df = 3; p = 0.9) (Figure 4F).

**Figure 4:**
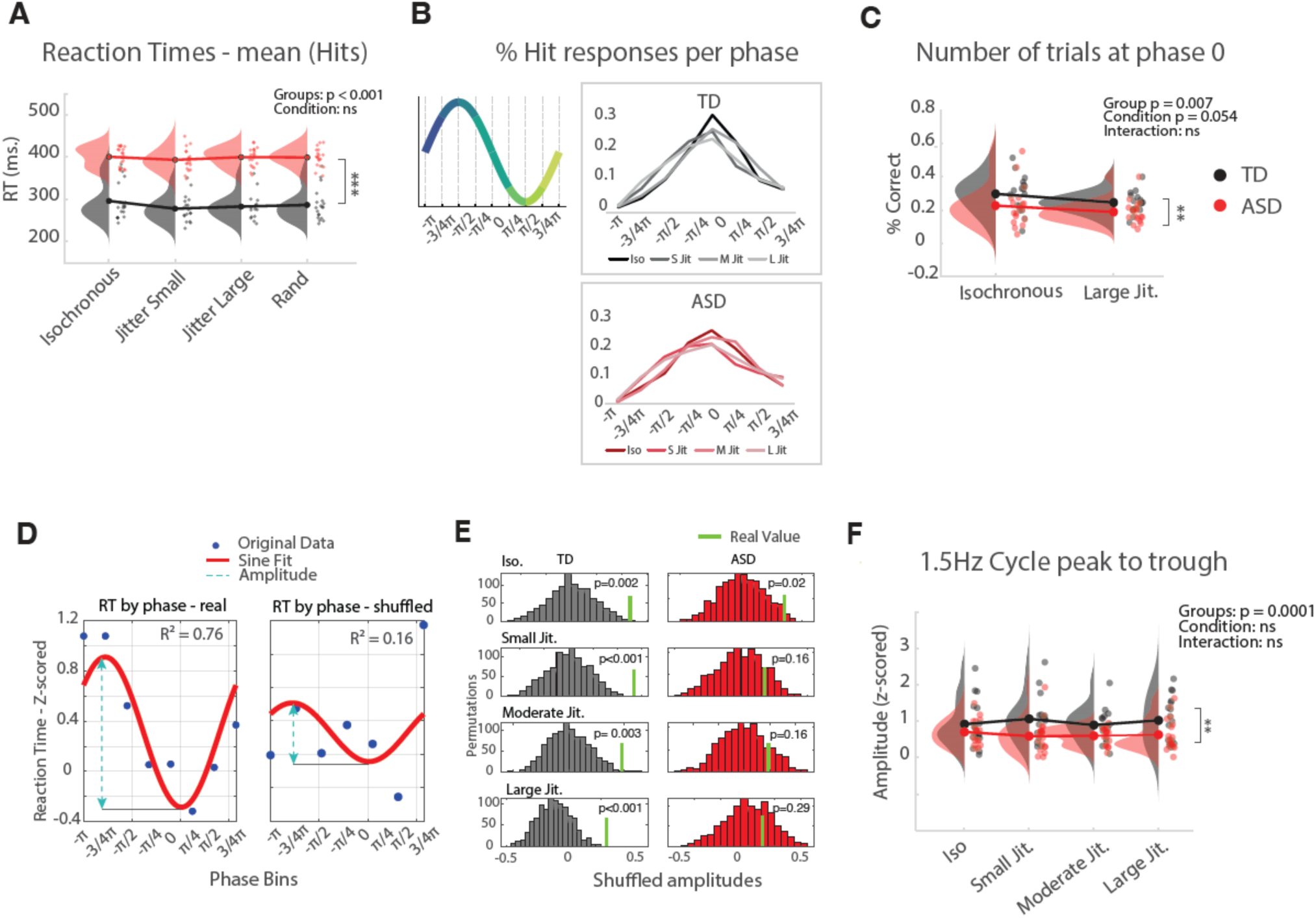
Task performance is slower across periodicities, but coupling with Delta oscillations is reduced in jitter conditions in the ASD group. **A.** Reaction times for group and condition show strong Group but not Condition effect or an interaction. **B.** Inset: illustration of one oscillation divided to 8 bins, as used in the analysis. Main: percentage of responses for different phases in the cycle. **C.** Stats for B: Main effects for response percentage were found for Group (*Isochronous* and *Random*) but only marginally for condition. **D**. Examples of sinusoid fitting. 1-cycle sinusoid was fitted for each participant to assess RT/phase coupling, for both real (left) and shuffled data (right). Peak-to-trough amplitudes were calculated for each sine fit. **E.** Permutation tests for each condition and group on real vs. shuffled peak-to-trough amplitudes show significant effect for all condition in NT group, but only in *Isochronous* condition in the ASD group. **F.** Peak-to-trough amplitudes show significant group effect, with lower amplitudes for ASD participants. Data in E and F is shown for fronto-central channels. (**p<0.01; *p<0.05).

This coupling is unique to the stimuli rhythm of 1.5Hz, since none of the other frequency bands we tested (theta, 5-7Hz; alpha, 9-11Hz) showed difference from zero, nor difference between TD or ASD (see Figure S2). The division of the RT to 8 bins yielded a low number of trials for some of the bins relative to others. To control for a possible effect of the different number of trials per bin, we pooled all trials from all participants and randomly chose the same number of trials from each bin (N=50), and repeated the analysis. We repeated this procedure 100 times, and averaged the sine fit across iterations. We observed the same pattern: Phase-RT coupling, indicated by the sine fit, was lower for ASD compared with TD group (TD: R^2^ = 0.85, 0.66, 0.77, 0.88; ASD: R^2^ = 0.58, 0.40, 0.66, 0.61, for *Isochronous*, *Jitter Small*, *Jitter Moderate* and *Jitter Large*, respectively). Rose plots for the target trials are seen in Figure S6.

### TRF model

The degree to which EEG activity in each of the regularity conditions can be predicted by the corresponding stimuli was evaluated using forward TRF models for each group and condition. The initial analysis was conducted using individual models, applied on each participant, focusing on a time lag of +500 to +2500 ms post stimulus onset, to specifically exclude the first evoked response from the analysis, on low-pass filtered (0.5–3 Hz) EEG. With this approach we aimed to assess predictive accuracy based on the ongoing rhythmic activity, rather than the initial evoked response, in low frequencies close to the frequency of stimulation. The model performance was assessed by calculating the correlation between predicted and actual EEG signals for a cluster of occipital electrodes (Figure 5). Group differences in model performance were analyzed using repeated measures non-parametric testing (Friedman’s ANOVA). As expected, prediction accuracy of the TRF model decreased with increasing temporal jitter, reflecting the reduced precision and reproducibility of stimulus encoding under less predictable conditions. A significant main effect was found (Friedman’s χ^2^ = 42.3; df = 7; p < 0.001). Post-hoc analyses (Durbin-Conover tests) revealed a significant between-group difference only for the *Small Jitter* condition (t = 2.62; p = 0.01), with the ASD group showing lower predictive accuracy compared to NT. There were no significant differences between groups for the *Isochronous* (t = 1.52; p = 0.13), *Moderate* (t = 1.27; p = 0.20), or *Large Jitter* conditions (t = 0.84; p = 0.39). To further assess model robustness, group-level models (i.e., subject-independent, where all sequences from all participants are pooled together) were trained with and without the first stimulus. This was applied to EEG in two filtering schemes: low-pass filtering (0.5-3Hz), as was done for the individual models, and high-passed filtering (5-30Hz), to evaluate the contribution of high-frequency components to the model’s predictive power. In the low frequencies, the group-level models showed significantly reduced prediction accuracy for the ASD group compared to NT in all four conditions (Figure S3), when the response to the first stimulus was included. However, when the first response was excluded from the EEG inputs to the model, the predictive power of the model dropped gradually for both groups, together with increasing noise levels. In addition, model predictions were lower for ASD than NT, in *Isochronous*, *Small Jitter* and *Moderate Jitter* conditions. When applied to higher frequencies (5-30 Hz), to evaluate the contribution of high-frequency components to the model’s predictive power, the prediction accuracy of the model dropped significantly (Pearson’s r<0.1) when the first stimulus was included, and even more (<0.01) when it was excluded, for the three jittered conditions in both groups (Figure S4). Overall, these findings imply that stimulus encoding in the EEG is predominantly driven by low-frequency components, with higher frequencies contributing primarily to the initial evoked response, and with generally a lower level of accuracy in the ASD, compared with the NT group.

**Figure 5:**
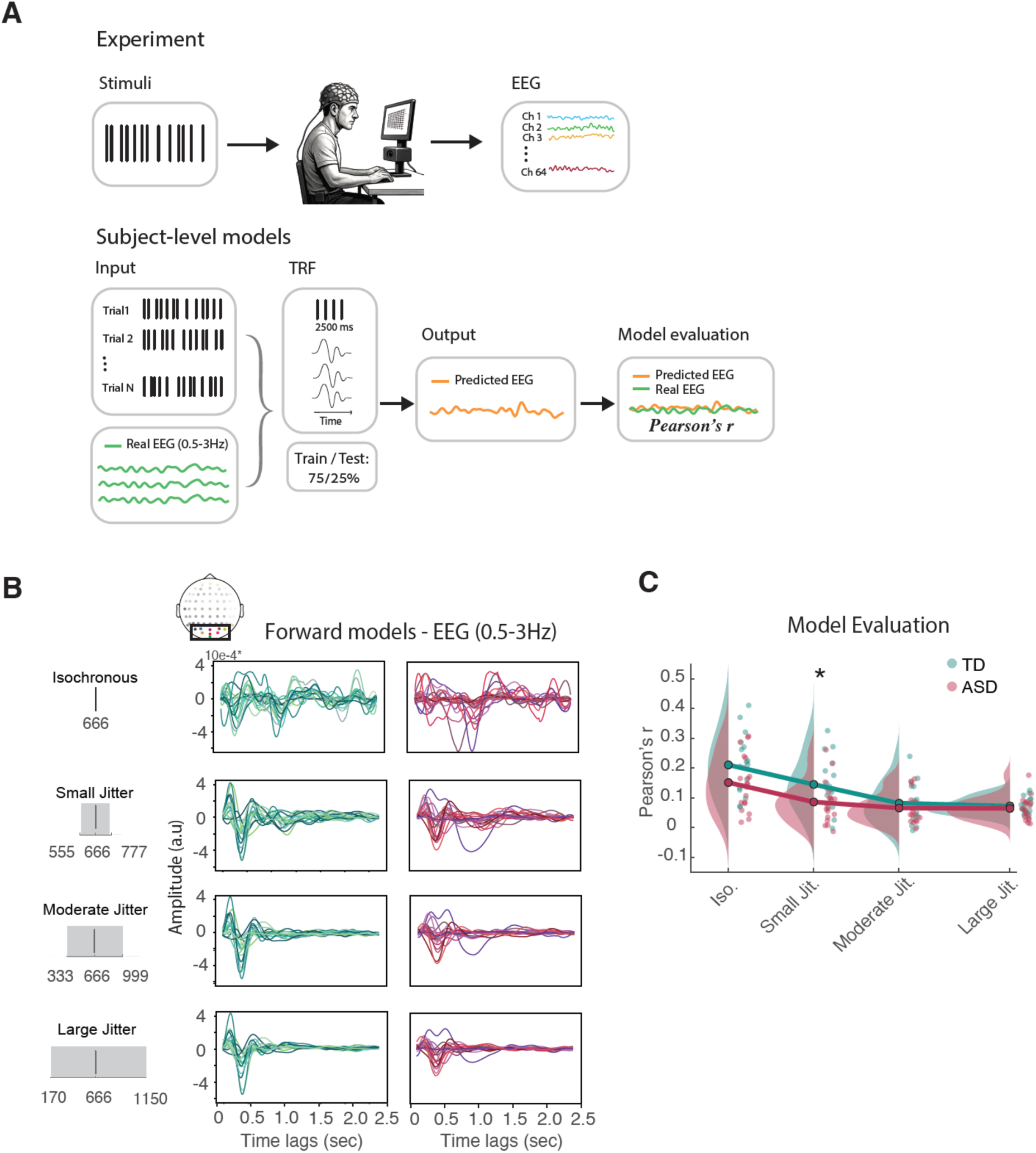
TRF forward model shows selective impairment in a Jitter condition for the ASD group. **A.** illustration of the procedure. **B.** TRF for each condition and group for an occipital cluster of electrodes (see Methods). Each line represents a TRF of one individual participant. **B.** Prediction accuracy measured as the Pearson correlation between model predicted EEG and real EEG per participant for time lags of 500-2500 ms. Results show significant group differences for Small Jitter condition.

### Sustained Entrainment

At silent mode, since the data was noisier when no stimulus is presented, we compared the two extreme periodic conditions: Isochronous vs. Random, at 1.5Hz (marked by the pink bar on Figure 6). We found a significant effect for Condition (F = 9.5; df = 1; p = 0.003), and a Group × Condition interaction (F = 4.37; df = 1; p=0.04), but no Group effect (F = 1.05; df = 1; p = 0.309). Post-hoc test shows condition effect within the TD group (p = 0.007).

**Figure 6:**
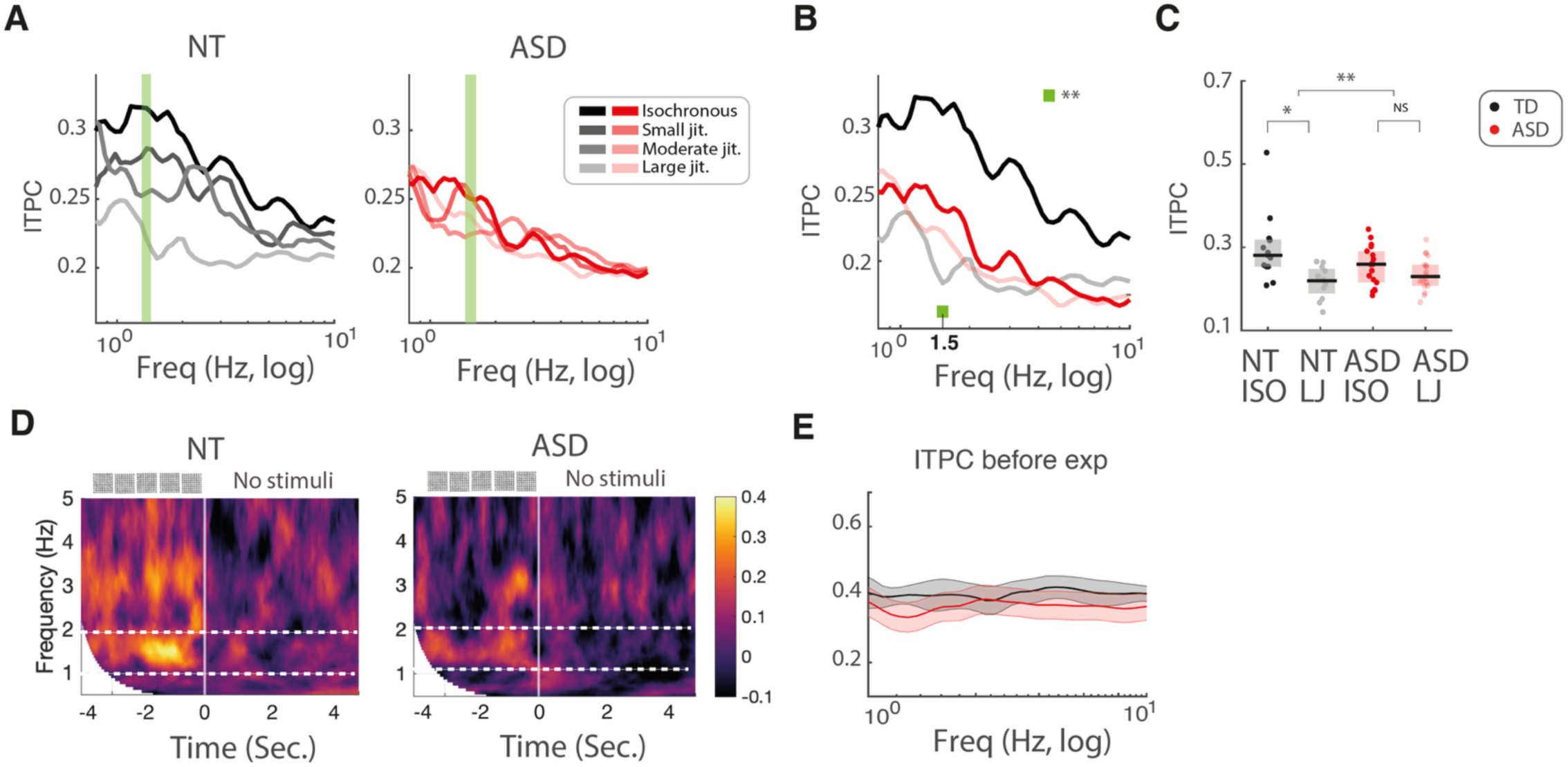
Sustained Entrainment. **A.** ITPC over 10 sec silent mode, following each of the 4 conditions, for NT and ASD. **B.** Isochronous and Random conditions are different for the groups at 1.5 Hz, showing significant Group x Condition interaction. **C.** Significant Group, and post-hoc effects. **D.** ITPC (*Isochronous*) before and after transition to silent mode. **E.** ITPC at silent mode before experiment, to control for baseline differences between the groups. *p < 0.05; **p < 0.01

### Clinical – Physiological Relationship

Correlation matrices were calculated for each group, combining physiological measures and clinical scores (See Figure 7). We were focused on correlations between physiological (or behavioral) measures and clinical scores. The ASD group shows strong correlation between full scale IQ (FSIQ) and ITPC modulation slope (*r = -0.51; p = 0.04, following FDR correction*), (see Figure 3D for ITPC slope), which denotes the strength of modulation to change in rhythmicity. The TD group shows significant negative correlation between performance measure of behavioral accuracy (Hit rate probability) and autism quotient (AQ) (*r = -0.58; p = 0.04, following FDR correction*) : the lower the performance, the higher the AQ.

**Figure 7:**
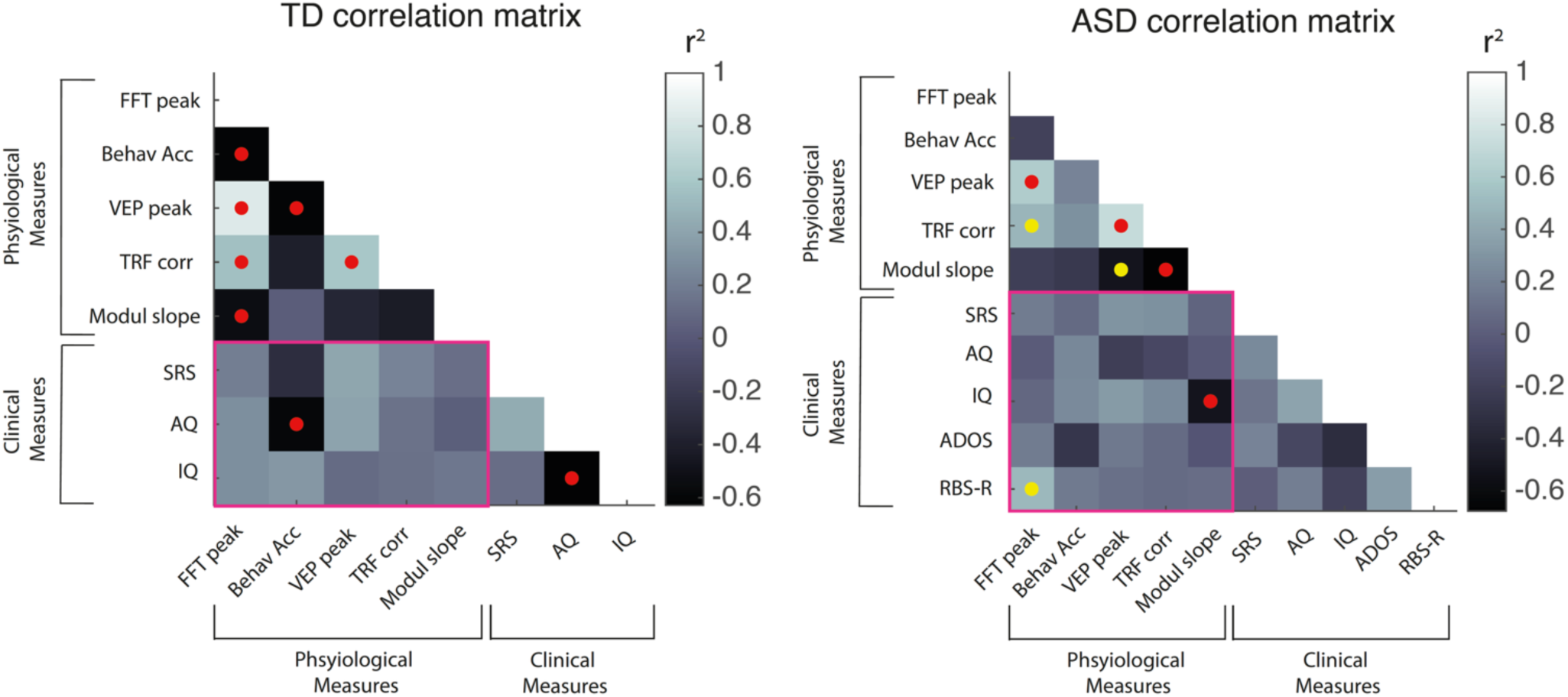
Correlation matrices for TD (left) and ASD shows significant EEG-clinical correlations. Red dots: Significant correlations that survived correction for false discovery rates (FDR). Yellow dots: correlations that were significant but did not survive FDR. Enclosed in the pink frame: correlations between physiological index and clinical scores. In the ASD group, modulation slope, indicating the ITPC modulation by the rhythmic conditions, is significantly correlated with IQ.

## Discussion

Reduced flexibility in processing temporal information is widely observed in autism spectrum disorder (ASD). This, and its impact on perception and performance, has been studied using behavioral (Chambon *et al*., 2017; Vishne *et al*., 2021; Patel *et al*., 2022), physiological (Thillay *et al*., 2016; Lawson *et al*., 2017a; Beker *et al*., 2021b; Cannon *et al*., 2023), and computational approaches (Lieder *et al*., 2019; Vishne *et al*., 2021; Wertheimer & Hart, 2024). Evidence converges on the claim that individuals with ASD show impairments in applying predictive rules, which affects their interaction with dynamic environments (Beker & Molholm, 2023), but they lack characterization of this impairment and how it modulates on regularity of sensory stimuli. We hypothesized that neural entrainment would be particularly impaired when flexible adaptation to jittered input is required. We systematically manipulated temporal regularity to test whether deficits in ASD reflect a general impairment or a specific vulnerability to noisy rhythms. If the results pointed to an overall impairment, regardless of the jitter level, this would indicate a general impairment in entraining to regularities. If, on the other hand, entrainment was more severely impaired when regularities exist but are noisy, it would indicate a more drastic impairment in jittered rhythms, where flexible processing of noisy conditions is demanded, but not met.

### Frequency analysis and ITPC modulation by regularity

Our results show a mixed pattern when it comes to supporting a general impairment in processing regularities vs. a greater vulnerability to volatile regularities. Power at the average frequency of stimulation (1.5Hz) showed a gradual effect of regularity level: power decreased as regularity decreased (Figure 2) and, while omnibus ANOVA did not reveal a significant Group x Condition interaction, a complementary condition-wise permutation tests provided a more focused analysis, revealing group differences in the Small and Moderate conditions: The ASD group had significantly lower 1.5Hz power than NT in *Small Jitter* and *Moderate Jitter*, but not in *Isochronous,* where both groups showed a similar power at 1.5Hz or *Large Jitter* condition, showing minimal or no power.

ITPC results support a general impairment, but also reduced modulation as regularity decreases: while both groups showed reduced neural entrainment as temporal regularity decreased, the ASD group exhibited a markedly attenuated modulation of this effect. ITPC showed a general reduction in phase-locking with increased irregularity across all participants, but this gradation was weaker in the ASD group, particularly in posterior scalp regions. Together, these findings suggest that individuals with ASD are less sensitive to changes in temporal regularity, displaying a blunted neural tracking of rhythmic structure. This reduced modulation may reflect atypical encoding of temporal predictions.

### TRF modeling

TRF models, which estimate how well neural responses track stimulus timing, mirrored these findings. TRF modeling complements traditional analyses of neural entrainment which capture oscillatory alignment but may not fully account for how neural responses encode stimulus dynamics over time. TRF explicitly models the temporal relationship between rhythmic input and brain activity, enabling us to assess how well the neural system tracks structured variability in the stimulus stream(Lalor *et al*., 2009; Crosse *et al*., 2021). We applied both individual-level models, where EEG signals for the training and testing sets are taken from the same individual, and group-level model, where the EEG signals from all participants are pooled together. When individual TRF models were applied, model performance was reduced in all three *Jitter* conditions in both groups, compared with the *Isochronous* condition. Additionally, model performance was poorer for the ASD compared to NT overall, with greatest difference in the *Small Jitter* condition. In the group-level model, stronger differences were revealed between ASD and NT, compared to subject-level models, highlighting the greater inter-individual variability within the ASD group and emphasizing the value of subject-specific modeling approaches over methods that rely on across-subject averaging.

### Phase-performance coupling

Phase entrainment in delta frequencies to anticipated stimulus is known to have effects on reaction times (Stefanics *et al*., 2010; Gray *et al*., 2015; Lakatos *et al*., 2019). However, the effect of phase entrainment on performance in ASD has not been reported. By measuring RT-phase coupling we tested if there were optimal phases for target detection in individuals with ASD, as shown for NTs with rhythmic visual stimuli (Wilson & Foxe, 2020), and how this preference is modulated by reduced regularities. When compared to shuffled data, where no coupling exists, RT-phase coupling in ASD was only significant in the Isochronous condition, where a full prediction of the stimulus could be made, and lead to a facilitated response. This contrasts with the NT group, where phase of the 1.5Hz oscillations governed reaction times across conditions, regardless of stimulus regularity. This general impairment in ASD with an enhanced effect in the fully-regular rhythm is compatible with previous findings of poor sensory-motor synchronization to rhythmic stimuli in ASD (Vishne *et al*., 2021; Kasten *et al*., 2023; Wertheimer & Hart, 2024). Importantly, neither group showed any evidence of RT–phase coupling at the other relevant frequencies (theta and alpha) under any condition. Theta activity is associated with memory (Herweg *et al*., 2020) and cognitive control (Senoussi *et al*., 2022), whereas alpha is linked, among other function, to visual attention and sampling (VanRullen, 2016). Lack of coupling with these frequencies suggests that the effect we found was specific to the stimulus-driven regularities, and not induced oscillations, or to a general property of oscillatory activity outside the stimulation frequency range. A caveat related to this is the clear preference of number of Hit response for the optimal phases, yielding a very few responses in the outlier bins (-pi, 3/4pi). To rule out the effect of different number of trials in the bins, we performed a control test where we pooled together all trials from all subjects within each group, while limiting number of trials to randomly permutated the same number of trials (N=50) from each bin and repeated the analysis. The same pattern, where the RT-phase coupling was higher for NT than the ASD group across condition, was observed here, suggesting that this relationship is robust.

### Sustained Entrainment

To find if the impact of rhythmic stimuli lasted beyond the presentation period, while ruling out the possible confounding of rhythmic entrainment with stimulus evoked response, we extended the EEG recording for 10 seconds immediately following the end of each condition block. We hypothesized that oscillatory activity during this “silent” period, when no visual stimulus was presented, would indicate a strong internal oscillatory entrainment to previously presented stimuli that was sustained when stimulation was over. We found that the modulation by the rhythmic conditions persisted in the NT participants even after the visual stimuli ceased. This was not the case in ASD group: *Isochronous* and *Jitter Large* conditions were comparable within the ASD group (Fig 6b). Furthermore, between group analysis revealed that the *Isochronous*–following ITPC was stronger in NT than in ASD (Fig. 6C).

### Correlation with clinical scores

We found a significant correlation between the modulation index of neural entrainment across regularity conditions and IQ within the ASD group. This relationship provides initial evidence that individual variability in sensitivity to temporal regularity may map onto core clinical features of ASD. While correlation with ADOS have not been revealed, this underscore the translational relevance of rhythmic entrainment as a candidate physiological marker, and point to its potential utility for stratifying individuals by clinical phenotype. By establishing this link, we provide evidence that disruptions in neural synchronization to environmental rhythms may underlie specific aspects of the behavioral profile in autism.

## Conclusions

Our results demonstrate reduced neural and behavioral entrainment to rhythmic stimuli in ASD, particularly when *Small* and/or *Moderate* temporal irregularity is introduced. This selective intolerance to jitter may reflect heightened sensory precision (O’Riordan *et al*., 2001; Bonnel *et al*., 2003a; Happe & Frith, 2006; Robertson *et al*., 2013), combined with reduced adaptability to environmental dynamics (Vishne *et al*., 2021; Beker *et al*., 2021b; Cannon *et al*., 2023), affecting the ability to anticipate events in variable, unstable environments, with potential consequences for adapting to sensory contexts. Although not definitive, reduced RT-phase coupling and lower TRF model performance support a diminished capacity to adapt to changing temporal structures.

Several theoretical frameworks align with our findings. The hypo-prior account proposes underweighting of prior information, while aberrant precision and predictive coding models implicate inflexible sensory weighting and high prediction error (Pellicano & Burr, 2012b; Lawson *et al*., 2017b). Although our results do not directly resolve these theories, they support a broader disruption in predictive processing.

A recent model (Wertheimer & Hart, 2024) suggests uniform allocation of encoding resources across stimulus intensities in ASD as an explanation to the poor input discrimination at the majority, central range of stimulus intensities, with a high discrimination at the extremes. Importantly, this model predicts uniform allocation of encoding resources, which is supported by our finding of reduced modulation of ITPC and Phase-RT by the rhythms. While the biology of jitter intolerance remains outside the scope of this study, prior work links altered neural synchrony to connectivity disruptions (Assaf *et al*., 2010; Weng *et al*., 2010; Di Martino *et al*., 2014; Long *et al*., 2016). Rhythmic entrainment, built upon highly precise timing, might be one of the first processes to be impaired and to show deficits in applying and weighing prior information when processing sensory inputs which require adaptation to regularities (Pellicano & Burr, 2012a; Van de Cruys *et al*., 2014; Lawson *et al*., 2017b).

## Limitations

The relatively low number of target stimuli in each of the phase bins, when testing the phase distribution in the RT-Phase coupling analysis limits our conclusions that can be drawn from this analysis. The reason for this low number is that target trials comprise 16% of all trials in the experiment, resulting in a relatively small number of trials in each of the eight phase bins.

## Supplementary Material

**Fig S1:**
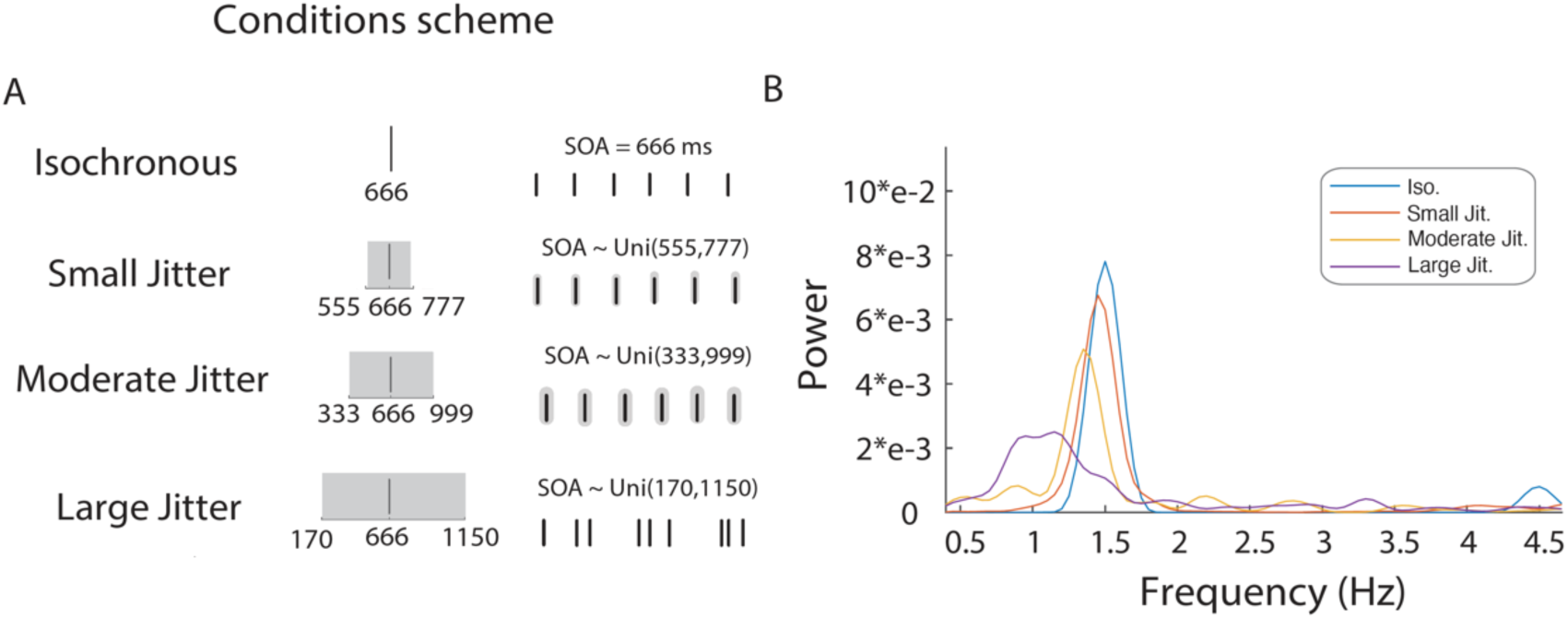
A. Schematic of condition for the different conditions. B. Spectral analysis of the stimuli, as recorded from the screen by photodiode.

**Figure S2.**
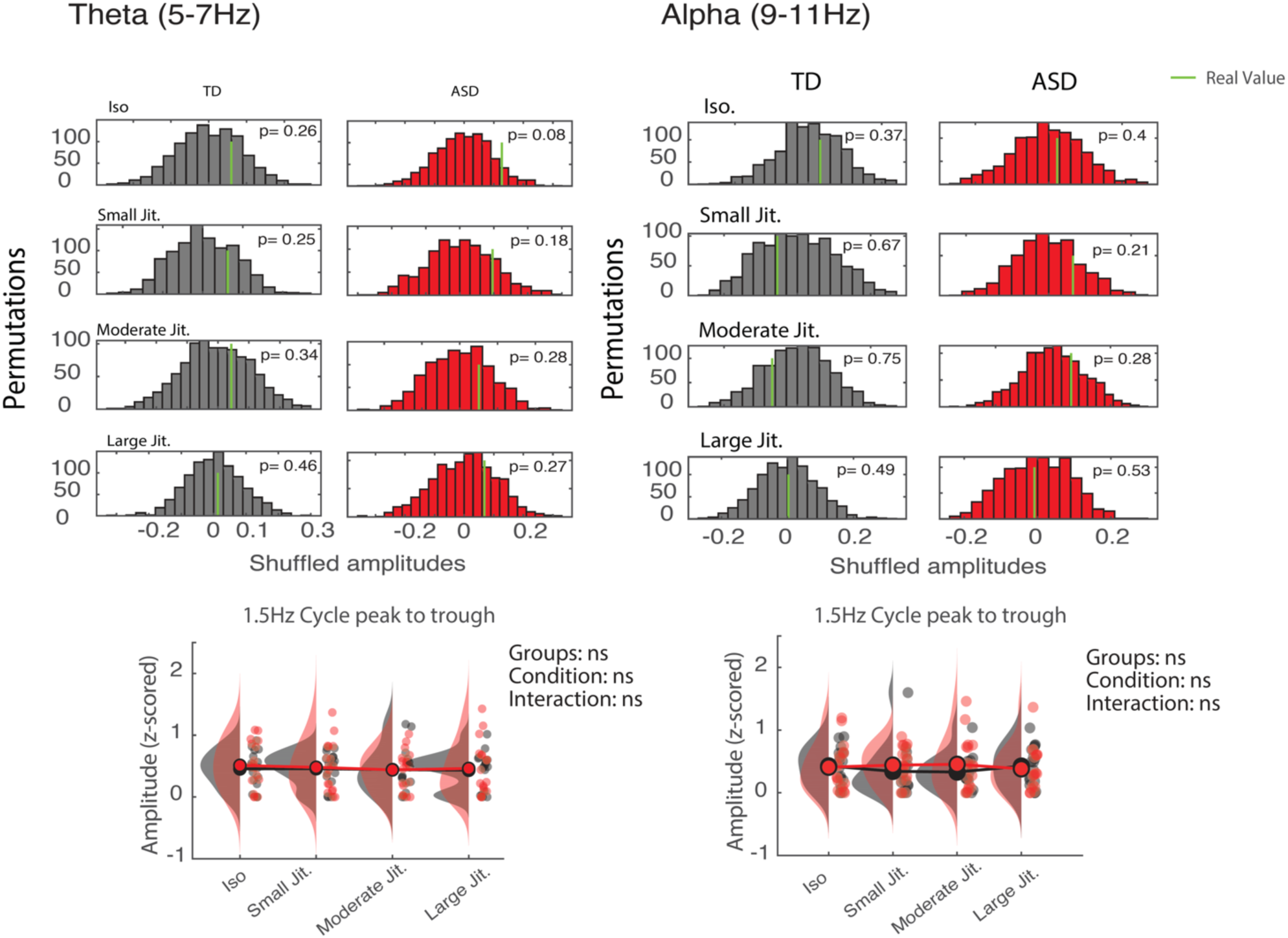
No RT-Phase coupling in any of the groups, or Group differences in Theta or Alpha frequencies.

**Figure S3:**
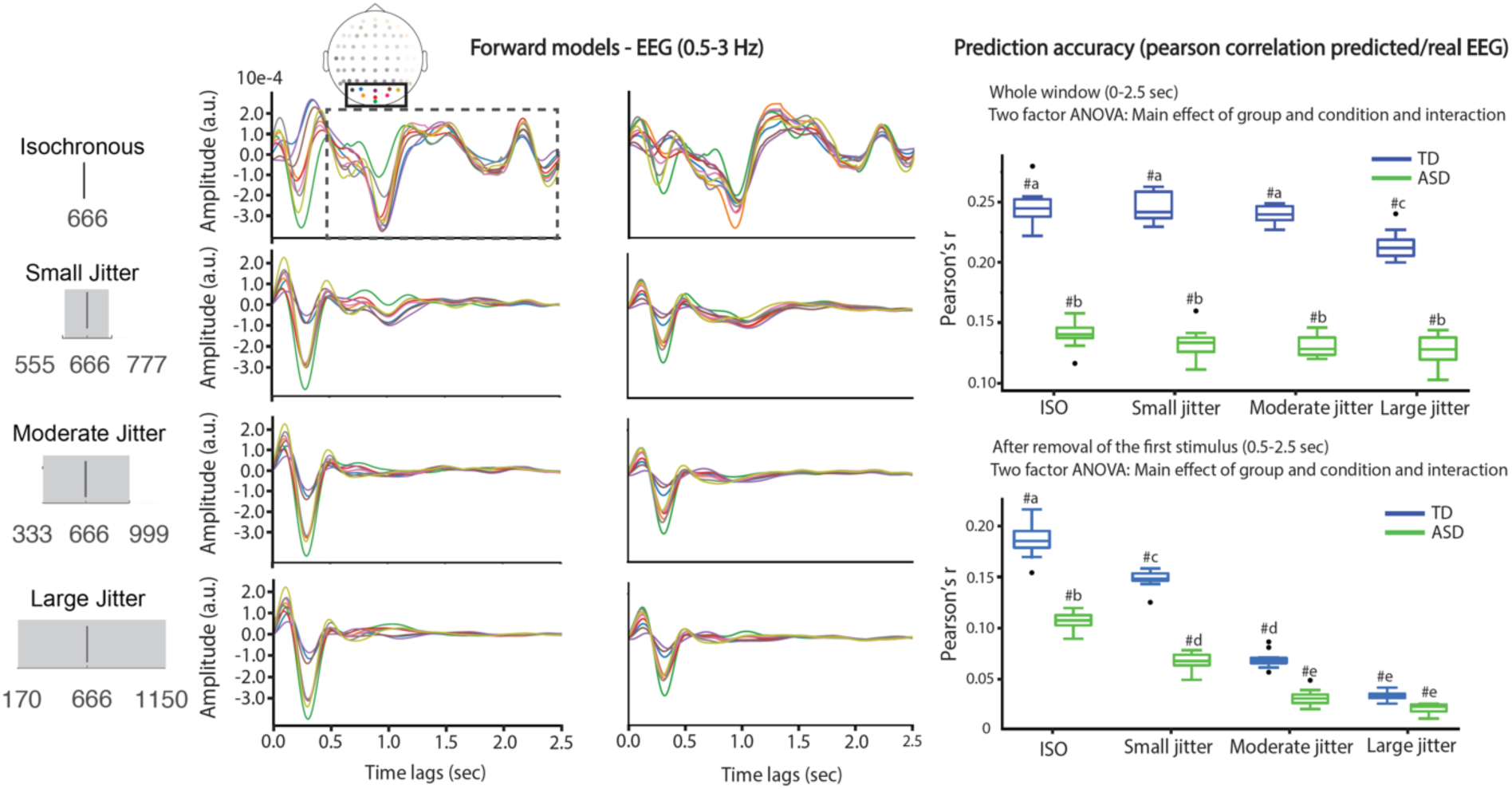
TRF forward model based on group models, low pass filtered.

**Figure S4:**
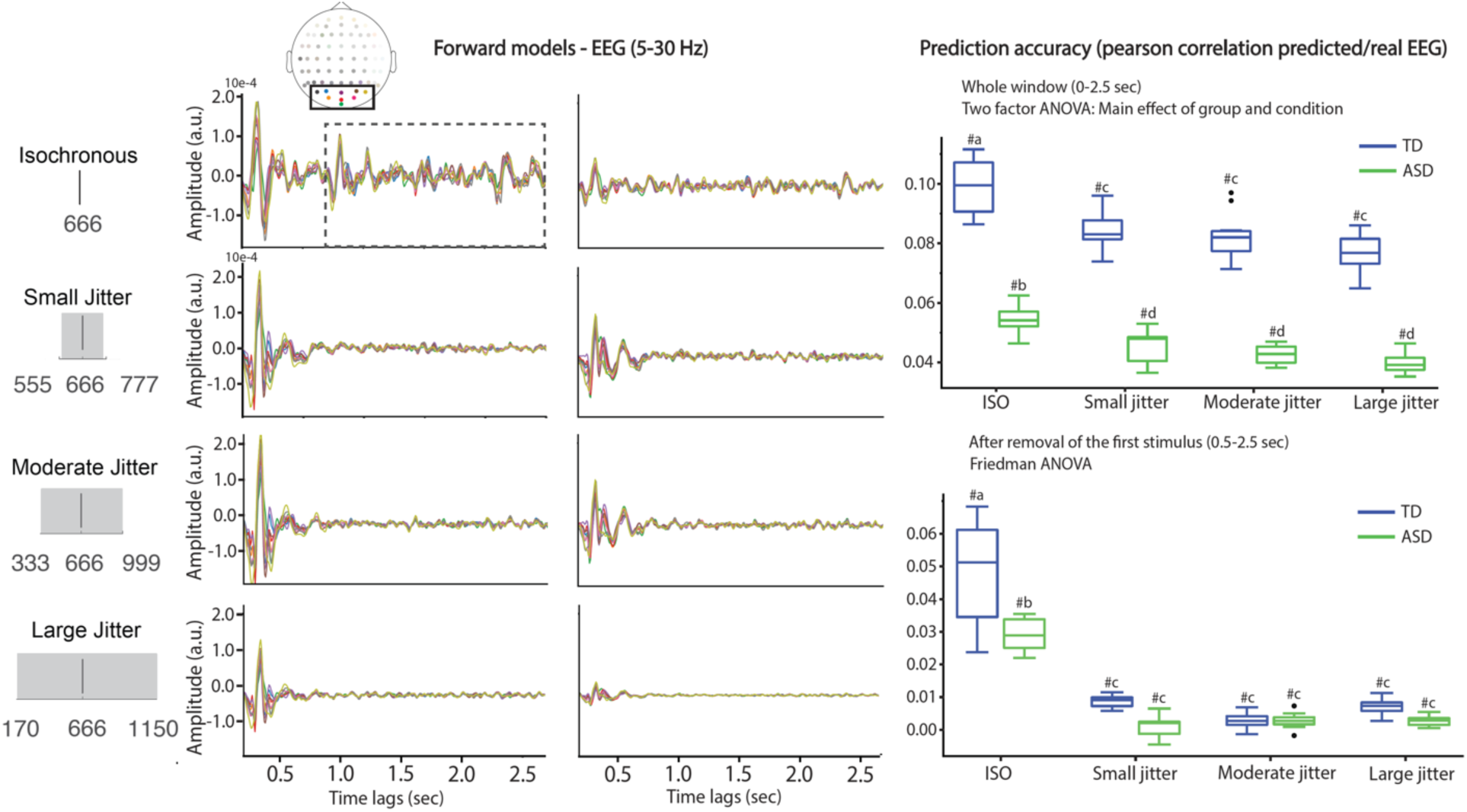
TRF model based on group models, high-pass filtered.

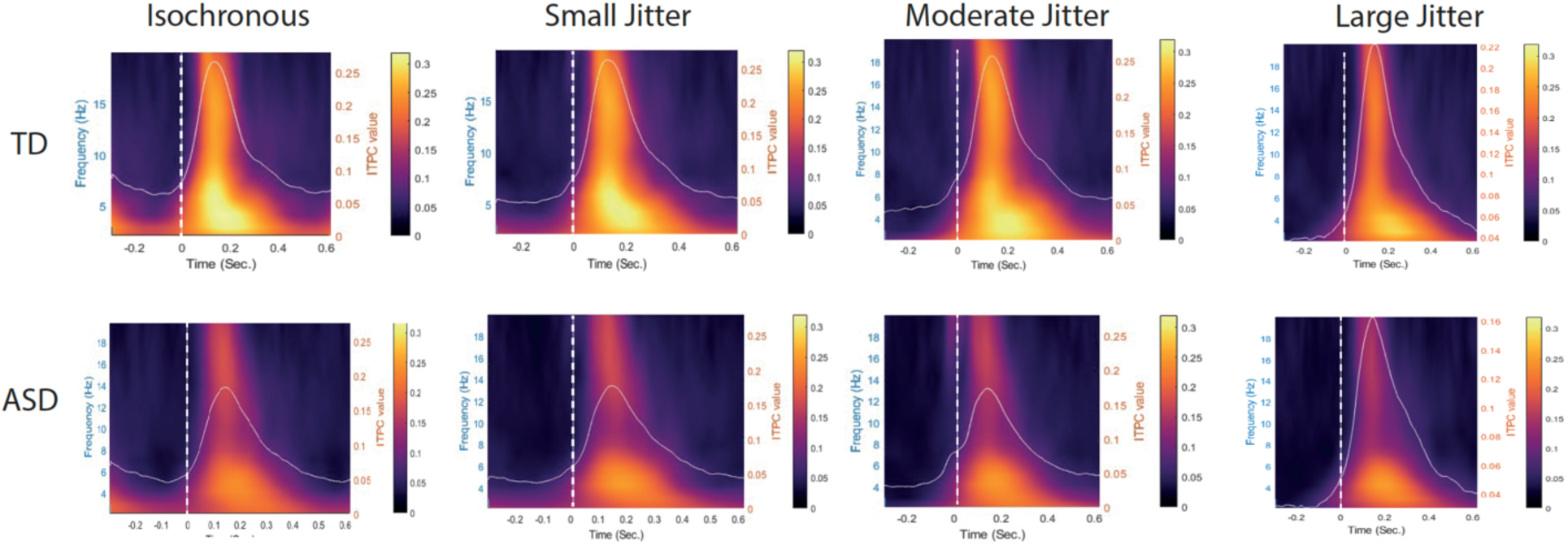

**Figure S6:**
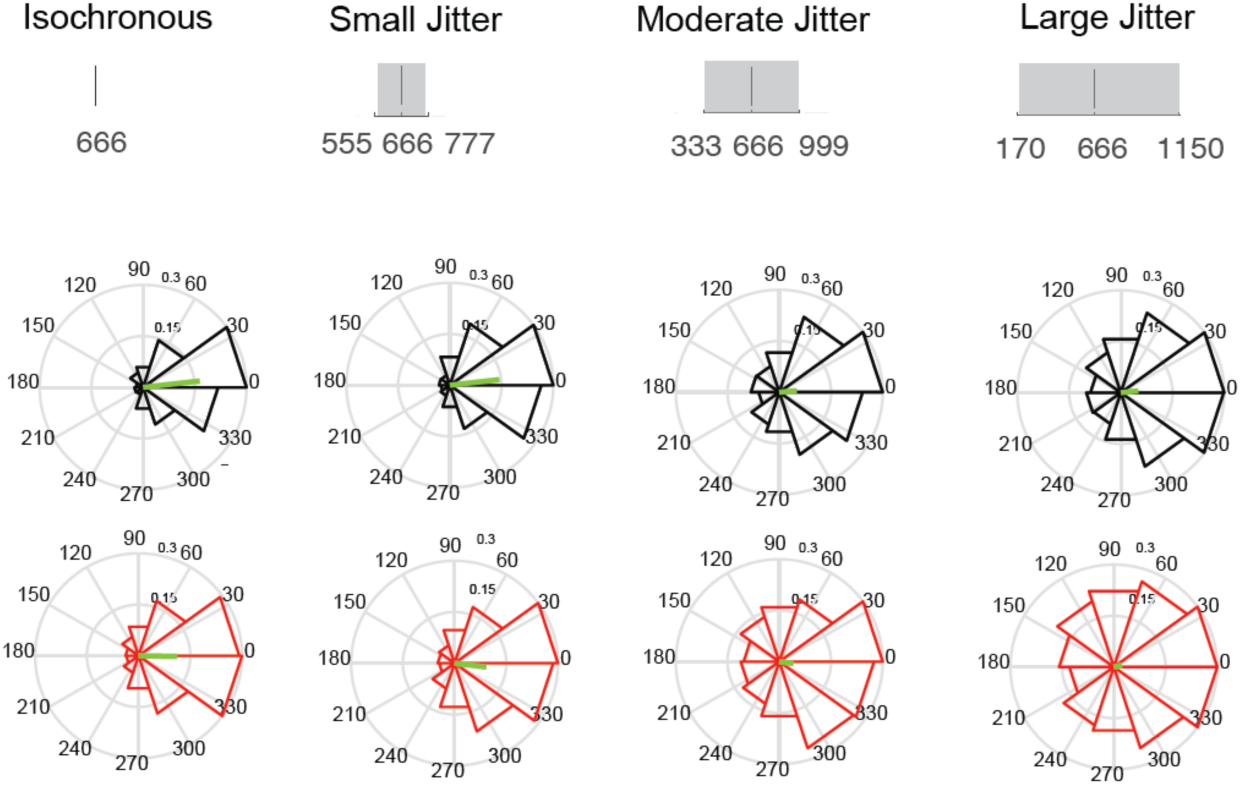
rose plots for phases during target presentation show phase concentration for each group and condition. Except ASD *Large Jitter* (p = 0.42), phase concentration was significant to all group/conditions (p <0.0001).

**Table S1:** Demographics (Mean±SD) for NT and ASD groups. Stimulation timing: Prior to analysis, we tested whether the tempos of the stimuli presented on the screen had the desired frequency content. For that, we used an oscilloscope to record the light emitted from the screen during stimuli presentation and received by a photodiode. We recorded one block from each condition. Time-frequency analysis was then performed on this time-series of voltage fluctuations and shows the fundamental 1.5Hz for the conditions, with a small reduction in frequency as tempo irregularity increased. This is due to the nature of SOA distribution, which yields a minor increase in the overall average SOA towards the end of the segment.

| Column1 | NT | ASD |
| --- | --- | --- |
| N | 21 (2 F) | 20 (10 F) |
| Age | 21.17±5.8 | 22.5±3.4 |
| IQ (Full scale) | 108±13.8 | 104±15.8 |
| ADOS score | N/A | 8.1±0.5 |

